# Multidimensional analysis and detection of informative features in diffusion MRI measurements of human white matter

**DOI:** 10.1101/2019.12.19.882928

**Authors:** Adam Richie-Halford, Jason Yeatman, Noah Simon, Ariel Rokem

**Affiliations:** Department of Physics, University of Washington, Seattle, WA, 98105, USA; Graduate School of Education and Division of Developmental and Behavioral Pediatrics, Stanford University, Stanford, CA, 94305, USA; Department of Biostatistics, University of Washington, Seattle, WA, 98105, USA; eScience Institute, University of Washington, Seattle, WA, 98105, USA

## Abstract

The white matter contains long-range connections between different brain regions and the organization of these connections holds important implications for brain function in health and disease. Tractometry uses diffusion-weighted magnetic resonance imaging (dMRI) data to quantify tissue properties (e.g. fractional anisotropy (FA), mean diffusivity (MD), etc.), along the trajectories of these connections [1]. Statistical inference from tractometry usually either (a) averages these quantities along the length of each bundle in each individual, or (b) performs analysis point-by-point along each bundle, with group comparisons or regression models computed separately for each point along every one of the bundles. These approaches are limited in their sensitivity, in the former case, or in their statistical power, in the latter. In the present work, we developed a method based on the sparse group lasso (SGL) [2] that takes into account tissue properties measured along all of the bundles, and selects informative features by enforcing sparsity, not only at the level of individual bundles, but also across the entire set of bundles and all of the measured tissue properties. The sparsity penalties for each of these constraints is identified using a nested cross-validation scheme that guards against over-fitting and simultaneously identifies the correct level of sparsity. We demonstrate the accuracy of the method in two settings: i) In a classification setting, patients with amyotrophic lateral sclerosis (ALS) are accurately distinguished from matched controls [3]. Furthermore, SGL automatically identifies FA in the corticospinal tract as important for this classification – correctly finding the parts of the white matter known to be affected by the disease. ii) In a regression setting, dMRI is used to accurately predict “brain age” [4, 5]. In this case, the weights are distributed throughout the white matter indicating that many different regions of the white matter change with development and contribute to the prediction of age. Thus, SGL makes it possible to leverage the multivariate relationship between diffusion properties measured along multiple bundles to make accurate predictions of subject characteristics while simultaneously discovering the most relevant features of the white matter for the characteristic of interest.

## Introduction

Diffusion-weighted Magnetic Resonance Imaging (dMRI) provides a unique view into the physical properties of the connections that comprise the brain white matter. While the measurements are usually conducted with voxels at the millimeter scale, water molecules within each voxel diffuse with characteristic lengths at the micrometer scale, providing aggregate information about the physical structure of the white matter, including the density of axons and distribution of fiber orientations within each voxel [6]. Even though metrics derived from diffusion measurements are ambiguous in terms of their underlying biological interpretation [7], analyzing the variance in these properties has proven useful in characterizing individual differences in cognitive function, characterizing differences between populations and detecting brain abnormalities associated with disease [8].

To relate the diffusion in each voxel to the macro-structure of long-range connections between different brain regions, methods for computational tract-tracing from diffusion MRI, or tractography, combine the estimates of fiber orientations in each voxel to form streamlines that traverse the volume of the white matter [9, 10]. These methods have been under increased scrutiny and several lines of investigation have raised critiques of their validity [11, 12]. On the other hand, there have been efforts to shore up the inferences made with these methods [13–18]. Importantly, though discovering novel tracts requires extraordinary evidence, and delineating the exact cortical termination of the streamlines in the gray matter is still prone to error, there is little dispute that tractography can accurately define the location of several major white matter tracts that are known to exist within the core of the white matter [11, 19].

Leveraging this fact, one of the most powerful methods currently available to put macro- and micro-structure together is *tractometry* : assessment of the physical properties of the white matter along specific tracts [20]. Though there are several different available implementations of this overall idea, the principles are similar [1, 21–23]: tractometry begins by delineating the parts of the white matter that belong to different major “tracts” (i.e. anatomical or functional groups of white matter fibers), such as the corticospinal tract or arcuate fasciculus, assigning tractography generated streamlines to “bundles,” which approximate the anatomical tracts, and sampling biophysical properties (such as fractional anisotropy or mean diffusivity) along the length of these bundles. the parts of the white matter that belong to different major tracts (i.e. anatomical or functional groups of white matter fibers), such as the corticospinal tract or arcuate fasciculus, assigning tractography generated streamlines to “bundles,” which approximate the anatomical tracts, and sampling biophysical properties (such as fractional anisotropy or mean diffusivity) along the length of these bundles. In some previous tractometry-based studies, tissue properties along the length of each tract were summarized by taking the mean along each bundle, but there is a large body of evidence showing that there is systematic variability in the values of diffusion metrics along the trajectory of each bundle. This justifies retaining the individual samples along the length of each bundle [1, 23, 24]. While this retains important information about each individual’s white matter, it also presents statistical challenges due to the dimensionality of the data. Based on tractometry, researchers may choose to compare different individuals to each other. This is usually done according to one of the following approaches:

1. Mass univariate approaches: In this approach comparisons between groups or across individuals are done independently at each node of each bundle, for each one of the diffusion metrics available at that point. This approach is exhaustive, but statistical power is compromised by a multiple comparison problem. Different approaches can be taken to resolving this challenge. For example, Colby and colleagues [24] used a non-parametric resmapling approach to correct for family-wise error across the different possible comparisons [25, 26].
2. Region of interest(ROI)-based approaches: An alternative that circumvents the multiple comparison problem is to select just a few tracts to compare in each individual, or even focusing on particular segments of these tracts based on *a priori* hypotheses. This approach is very powerful when the biological basis of the process of interest is relatively well understood (for a recent example, see [27]).
3. ROI-based selection, followed by multivariate analysis: Here, an ROI is selected based on *a priori* knowledge, and all the nodes or voxels in the ROI are used together to fit a model that can predict differences between individuals. An example of that is the “profilometry” framework, in which different diffusion metrics from a single tract are combined together to provide input to a multivariate analysis of covariance, and linear discriminant analysis [28].

Generally speaking, analysis methods should balance predictive accuracy with descriptive power [29, 30]. Accordingly, tractometry analysis should simultaneously capitalize on all the data across all tracts to make the best possible prediction, while also retaining and elucidating spatial information about the locations that are most informative for a prediction. In the present work, we developed a novel framework for analysis of tractometry that simultaneously selects the features for analysis, and fits a model to these features. We use a linear modeling approach, which aims to predict phenotypical variance in a group of subjects, based on a linear combination of the features estimated with tractometry.

Using this approach, we first need to deal with the large and asymmetric dimensionality of the data: tractometry data usually has many more features (i.e., number of measurements per individual) than samples (number of subjects), which makes inferences from the data about phenotypical differences between individuals ill-posed. This regime is the target of several statistical learning techniques, and is often solved by various forms of regularization. For example, Tikhonov regularization shrinks the solution such that the sum of squared contributions from the individual features are minimized [31]. Another solution to the problem is provided by the Lasso algorithm, which instead minimizes the sum of the absolute values of contributions of each feature [32]. This tends to shrink to zero the contributions of many of the features, providing results that are both accurate and interpretable. When additional structure is available in the organization of the data, regularization algorithms can take advantage of this structure. For example, if the features lend themselves to a natural division into different groups, the group lasso (GL) can be used to select groups of features, rather than individual features [33]. The Sparse Group Lasso (SGL) elaborates on this idea by providing control both of group sparsity, as well as overall sparsity of the solutions [34]. Because the features measured with tractomery lend themselves to grouping based on the tracts from which each measurement is taken, GL and SGL could provide a useful tool for linear model fitting in problems of this form. Here we, first, develop an implementation of SGL that is well suited to the analysis of tractometry data and, second, demonstrate the power and flexibility of this approach by applying it to both classification (disease diagnosis) and continuous prediction (age) problems from previously published studies [3, 4].

## Materials and methods

### Data

Two different previously-published datasets were used here:

1. Diffusion MRI from a previous study of the corticospinal tract (CST) in patients with amyotrophic lateral sclerosis (ALS [3]), containing data from 24 ALS patients and 24 demographically matched healthy controls. These data were measured in a GE Discovery 750 3T MRI scanner at the Institute of Bioimaging and Molecular Physiology in Catanzaro. Informed consent was provided as approved by the Ethical Committee of the University “Magna Graecia” of Catanzaro. Voxel resolution was 2 × 2 × 2 mm^3^ and 27 non-colinear directions were measured with a 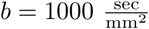. Data was preprocessed to correct for subject motion and for eddy currents. The diffusion tensor model [35] was fit in every voxel. We will refer to this dataset as ALS.
2. Diffusion MRI data from a previous study of properties of the white matter across the lifespan [4], containing dMRI data from 76 subjects with ages 6-50. These data were measured in a GE Discovery 750 3T MRI scanner at the Stanford Center for Cognitive and Neurobiological Imaging. The Stanford University IRB approved the procedures of this study. Informed consent was obtained from each adult participant, and assent for participation was provided by parents/guardians for children. Voxel resolution was 2 × 2 × 2mm^3^ with 96 non-colinear directions measured with a 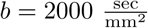 and 30 non-colinear directions measured with a 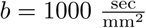. These data were acquired using a dual spin echo sequence, in which there is sufficient time for eddy currents to subside between the application of the gradients and the image acquisition, so no eddy current correction was applied, but motion correction was applied before fitting the diffusion tensor model [35] in every voxel using a robust fit [36]. We will refer to this dataset as WH.

Data from both of these studies was processed in a similar manner, using the Matlab Automated Fiber Quantification toolbox (AFQ) [1]: streamlines representing fascicles of white matter tracts were generated using a determinstic tractography algorithm that follows the prinicpal diffusion direction of the diffusion tensor in each voxel (STT) [37]. Major tracts were identified using multiple criteria: streamlines are selected as candidates for inclusion in a bundle of streamlines that represents a tract if they pass through known inclusion ROIs and do not pass through exclusion ROIs [38]. In addition, a probabilistic atlas is used to exclude streamlines which are unlikely to be part of a tract [39]. Each streamline is resampled to 100 nodes and the robust mean at each location is calculated by estimating the 3D covariance of the location of each node and excluding streamlines that are more than 5 standard deviations from the mean location in any node. Finally, a bundle profile of tissue properties in each bundle was created by interpolating the value of MRI maps of these tissue properties to the location of the nodes of the resampled streamlines designated to each bundle. In each of 100 nodes, the values are summed across streamlines, weighting the contribution of each streamline by the inverse of the mahalnobis distance of the node from the average of that node across streamlines. This means that streamlines that are more representative of the tract contribute more to the bundle profile, relative to streamlines that are on the edge of the tract.

This process creates bundle profiles, in which diffusion measures are quantified and averaged along twenty major fiber tracts. Here, we use only the mean diffusivity (MD) and the fractional anisotropy (FA) of the diffusion tensor, but additional dMRI-based maps or maps based on other quantitative MRI measurements can also be used. These bundle profiles, along with the phenotypical data we wish to explain or predict, form the input to the SGL algorithm. In a domain-agnostic machine learning context, the phenotypical data comprise the target variables while the bundle profiles form the feature or predictor variables (See Fig 1).

**Fig 1.**
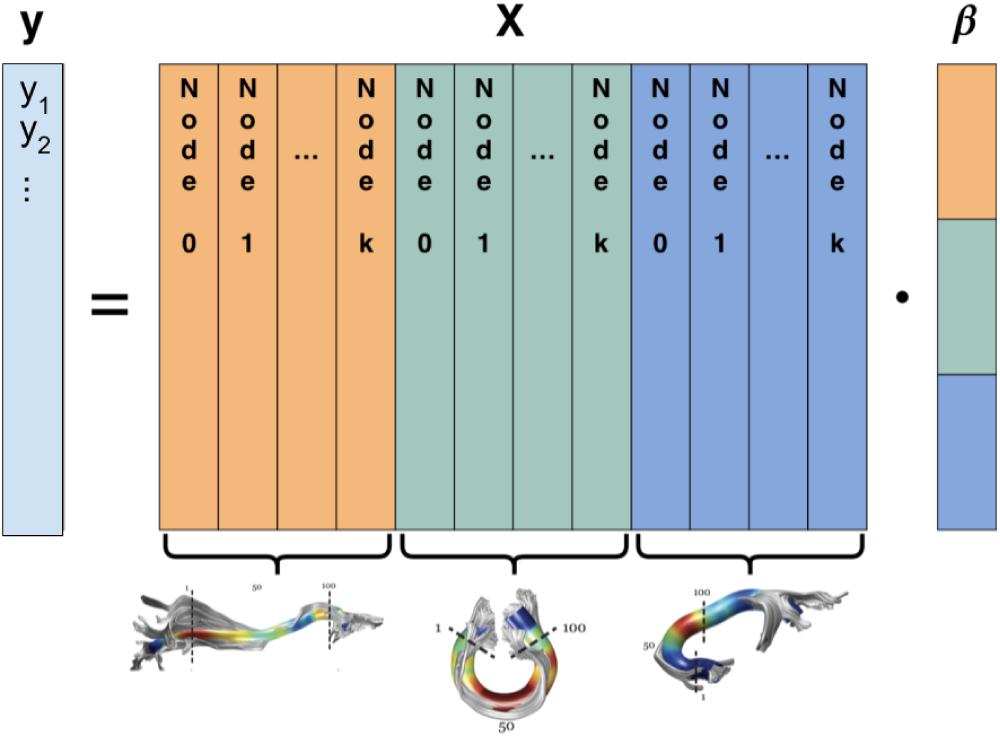
dMRI group structure. The phenotypical target data and tractometric features can be organized into a linear model, 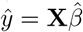. The feature matrix **X** is color-coded to reveal a natural group structure: the left (orange) group contains *k* features from the inferior fronto-occipital fascicle (IFOF), the middle (green) group contains *k* features from the corpus callosum, and the right (blue) group contains *k* features from the uncinate. The coefficients in 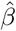 follow the same natural grouping. Fascicle image reproduced with permission from Ref [1] Figure 1.

### Sparse Group Lasso

Before fitting a model to the data, imputation and standardization are performed. Missing node values (e.g., in cases where AFQ designates a node as not-a-number) are imputed via linear interpolation. Care is taken not to interpolate across the boundaries between different bundles. Some diffusion metrics will have naturally larger variance than others and may therefore dominate the objective function and make the SGL estimator unable to learn from the lower variance metrics. For example, fractional anisotropy (FA) is bounded between zero and one and could be overwhelmed by an unscaled higher-variance metric like the mean diffusivity (MD). To prevent this we remove each feature’s mean and scale it to unit variance (z-score) using the StandardScaler from scikit-learn [40]. Scaling is performed separately within each cross-validation set’s training or testing data to prevent leakage of information between the testing and training sets [41].

After scaling and imputation, the tractometry data and target phenotypical data can be organized in a linear model:

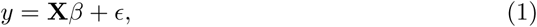

where *y* is the phenotype – categorical, such as a clinical diagnosis, or continuous numerical, such as the subject’s age. The tractometry data is represented in the feature matrix **X**, with rows corresponding to different subjects, and columns corresponding to diffusion measures at different nodes within each bundle. The relationship between tractometric features and the phenotypic target is characterized by the coefficients in *β*. The error term, *E* is an unobserved random variable that captures the error in the model. We denote our prediction of the targer phenotype as 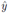 and the coefficients that produce this prediction as 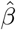, so that

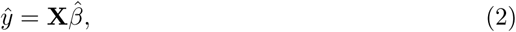

without the error term, *E*. In general, the feature matrix **X** has dimensions *S* × (*B* × *N* × *M*), where *S* is the number of subjects, *B* is the number of white matter bundles, *N* is the number of nodes in each bundle, and *M* is the number of diffusion metrics calculated at each node. Typically, *B* = 20, *N* = 100, and 2 ≤ *M* ≤ 8, resulting in ∼4, 000 −16, 000 features. Conversely, many dMRI studies have between a few dozen and a few hundred subjects, yielding a feature matrix that is wide and short. Even in cases where more than a thousand subjects are measured (e.g., in the Human Connectome Project, where 1,200 subjects were measured [42]), the problem is ill-posed: the high dimensionality of this data requires regularization to avoid overfitting and generate interpretable results.

Here, we propose that in addition to regularizing the coefficients in 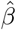, we can also capitalize on our knowledge of the group structure of the bundle profile features in **X**. The bundle-metric combinations form a natural grouping. For example, we expect that MD features within the left arcuate fasciculus will co-vary across individuals. Likewise for FA values within the right corticospinal tract (CST) and so on. This group structure is represented in Fig 1, which depicts the linear model 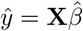. Thus, we seek a regularization approach that will fit a linear model with anatomically-grouped covariates, where we expect to observe both groupwise sparsity, where the number of groups (bundle/metric combinations) with at least one non-zero coefficients is small, as well as within-group sparsity, where the number of non-zero coefficients within each non-zero group is small. The sparse group lasso (SGL) is a penalized regression technique that satisfies exactly these criteria [2]. It solves for a coefficient vector 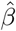 that satisfies

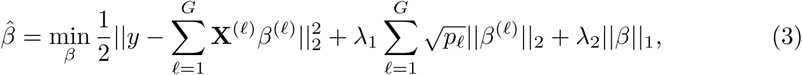

where *G* is the number of groups **X**^(*ℓ*)^ is the submatrix of **X** corresponding to group *ℓ, β*^(*ℓ*)^ is the coefficient vector for group *ℓ* and *p*_*ℓ*_ is the length of *β*^(*ℓ*)^. In the tractomtetry setting, *G* = *T* × *M* and ∀*ℓ* : *p*_*ℓ*_ = 100. The first term is the mean square error loss, *L*_mse_, as in the standard linear regression framework. The second and third terms encourage groupwise sparsity and overall sparsity, respectively. If *λ*_1_ = 0 and *λ*_2_ = 1, the SGL reduces to the traditional lasso [43]. Conversely, if *λ*_1_ = 1 and *λ*_2_ = 0, the SGL reduces to the group lasso [44].

### Incorporating transformations on *y*

Often, the target variable *y* is not in the domain in which the linear model can be best fit to it. Equation (2) can be slightly modified as:

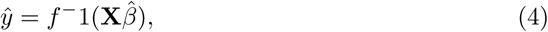

where the transformation function *f* ^*−*1^ characterizes the transform applied to the data before fitting the linear coefficients. For example, an often-used transform is the logarithmic transform:

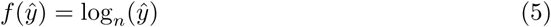

In this case, the model is parametrized by one additional fit parameter, *n*.

### Classification of categorical *y*

When the phenotypical target variable is categorical, as in the case of explaining or predicting the presence of a clinical diagnosis, the SGL is readily adapted to logistic regression, where the probability of a target variable belonging to an arbitrary defined “true” class is the logistic function of the result of the linear model,

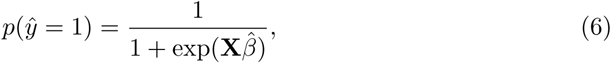

or equivalently, the mean squared error loss function in Eq (3) is replaced with the cross-entropy loss, which for binary classification is the negative log likelihood of the SGL classifier giving the “true” label:

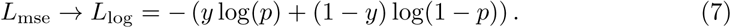

### Implementation, cross-validation and metaparameter optimization

For given values of *λ*_1_ and *λ*_2_, the cost function in Eq (3) can be optimized using proximal gradient descent methods [45] here implemented as a custom proximal operator that is then optimized using the C-OPT library [46]. This supplies an estimate of the optimal 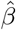 given a particular set of values for the meta-parameters *λ*_1_ and *λ*_2_.

To objectively evaluate the model and guard against over-fitting, we used a nested cross-validation scheme, depicted in Fig 2 for the categorical classification case. The subjects (i.e. rows of the feature matrix **X** in Fig 1 and Eq (1)) are shuffled and then decomposed into *k* batches, hereafter referred to as folds. For the ALS dataset we used *k* = 10 and for the WH dataset *k* = 5. For each unique fold, we hold that fold out as the test_outer_ set and let the remaining data comprise the train_outer_ set, with the subscript indicating the depth of the nested decomposition. We further decompose each train_outer_ set into three folds, and again for each unique fold, we hold out that fold as the test_inner_ set and let the remaining data comprise the train_inner_ set. At level-1 of the decomposition, we fit an SGL model using fixed regularization meta-parameters *λ*_1_ and *λ*_2_, training the model using train_inner_ and evaluating the fit on test_inner_. We find the optimal values for *λ*_1_ and *λ*_2_ using hyperoptimization, implemented using the hyperopt library’s `fmin` function [47] with a tree of Parzen estimators search algorithm [48]. For continuous numerical *y*, `fmin` searches for meta-parameter values that minimize the median absolute error. This can also be done minimizing the root of the mean squared error (RMSE) or to maximizing the coefficient of determination (*R*^2^). For binary categorical *y* `fmin` seeks to maximize the classification accuracy. This can also be done maximizing the area under the receiver operating curve (ROC AUC) or the average precision. Using hyperoptimization, we find optimal regularization parameters and 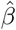 for each train_outer_ set and then use those to predict values for data in test_outer_. Thus each subject in the dataset has a predicted phenotype derived from a model that never saw its particular subject’s data.

**Fig 2.**
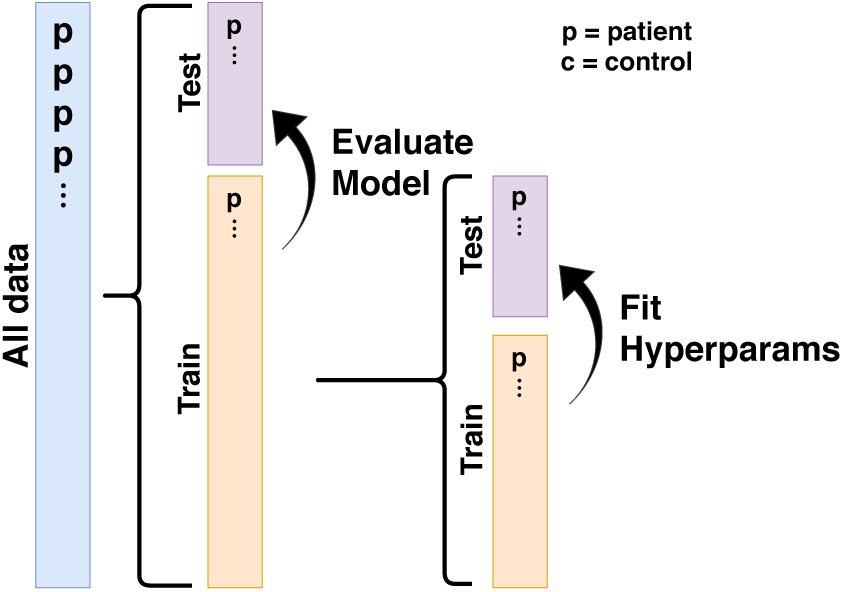
Nested *k*-fold cross-validation scheme. We evaluate model quality using a nested *k*-fold cross validation scheme. At level-0, the input data is decomposed three times into *k* shuffled groups and optimal hyperparameters are found for the level-0 training set. Optimization of these hyperparameters requires the use of the hyperopt library and many repeated evaluations of an SGL model over a search space of possible regularization parameters. These evaluations take place at level-1 of the decomposition, where the level-0 training set is further decomposed into three shuffled groups. For the ALS data, *k* = 10. For the WH data, *k* = 5.

The above procedure describes *k*-fold cross validation. In fact, we use repeated *k*-fold cross validation on the outer level of the decomposition, so that the input data is decomposed into *k* folds, three times. Thus, each subject has three predicted phenotypes. We then take the mean predicted value to summarize the performance of the model. In the classification case, the shuffling into folds is stratified such that each fold has a population that preserves the percentage of each class found in the larger input data.

### Software implementation

The full software implementation of the SGL approach presented here is available in a Python software package called AFQ-Insight, which is developed publicly in https://github.com/richford/afq-insight. The version of the code used to produce the results herein is also available in https://doi.org/10.5281/zenodo.3585942. AFQ-Insight reads the target and feature data that has been processed by AFQ from comma separated value (CSV) files conforming to the AFQ-Browser data format [49] and represents them internally as DataFrame objects from the pandas Python library [50]. The software provides different options for imputing missing data in the feature matrix. Missing interior nodes are imputed using linear interpolation. For missing exterior nodes, the user may choose between linear extrapolation and constant forward(back)-fill. Imputation uses only values from adjacent nodes in the same white matter bundle in the same subject so there is no danger of data leakage from other subjects. It uses the scikit-learn [40] library to decompose input data into separate test and train datasets, to scale each feature to have zero mean and unit variance, and to evaluate each fit in the hyperparameter search using appropriate classification and regression metrics such as accuracy, area under the receiver operating curve (AUC ROC), and coefficient of determination (*R*^2^). For each set of hyperparameters, we solve the SGL using a custom proximal operator supplied to the C-OPT library [46]. Appropriate hyperparameters are found using the hyperopt library [47].

## Results and discussion

We developed a method for analyzing dMRI tractometry data with SGL. We demonstrate the use of this method on two previously published datasets in both a classification setting and a regression setting.

### SGL accurately detects ALS in tractometry data in a classification setting

Using data from a previous study of the corticospinal tract (CST) profile and ALS [3], we tested the performance of SGL in a classification setting. The previous study predicted ALS status with a mean accuracy of 80% using a random forest algorithm based on a priori selection of features within the corticospinal tract. SGL delivers competitive predictive performance (mean 93% ± 2% accuracy, 0.978 ±0.006 ROC AUC) without the need for a priori feature engineering. The results of the classification prediction are shown in Figure 3 with “ground-truth” ALS status separated into columns, and predicted ALS status encoded by color. In addition to this classification performance, SGL also identifies the white matter tracts most important for ALS classification. The relative importance of white matter features is captured in the *β* coefficients from Eq (3). Figure 4 depicts these coefficients along the right CST, plotted over the FA values for the control and ALS subject groups. We find that SGL selects FA metrics in the corticospinal tract and particularly in the right corticospinal tract as most important to ALS classification, confirming previous findings [51–60] and identifying the portions of the brain that were selected *a priori* in the previous study from which we collected our data [3].

**Fig 3.**
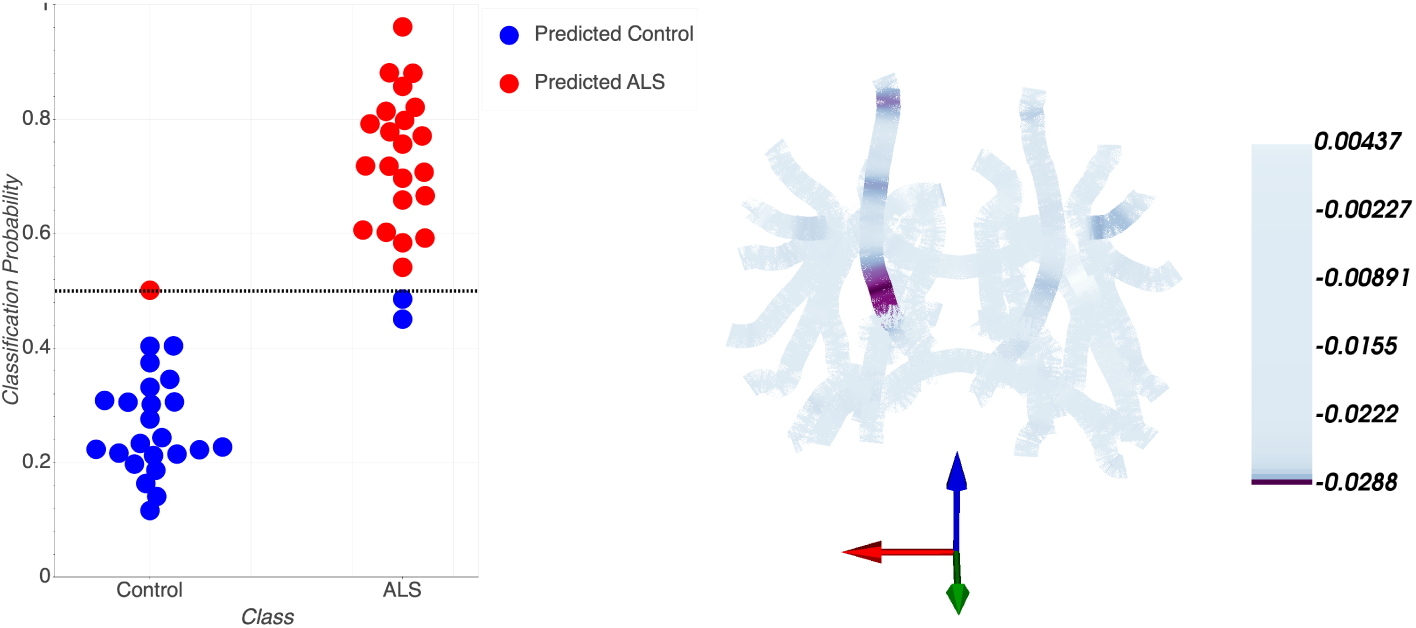
SGL accurately predicts ALS. Left: Classification probabilities for each subject’s ALS diagnosis. Controls are on the left while patients are on the right. Predicted controls are in blue and predicted patients are in red. Thus, false positive are represented as red dots on the left, while false negatives are represented as blue dots on the right. The SGL algorithm achieves 93% ± 2% accuracy, with 0.978 ±0.006 ROC AUC. Right: SGL coefficients are presented on a skeleton of the major tracts. The brain is oriented with the right hemisphere to our left and anterior out of the page. As expected large negative coefficients are in the FA of the CST (and particularly in the right hemisphere, here to the left)

**Fig 4.**
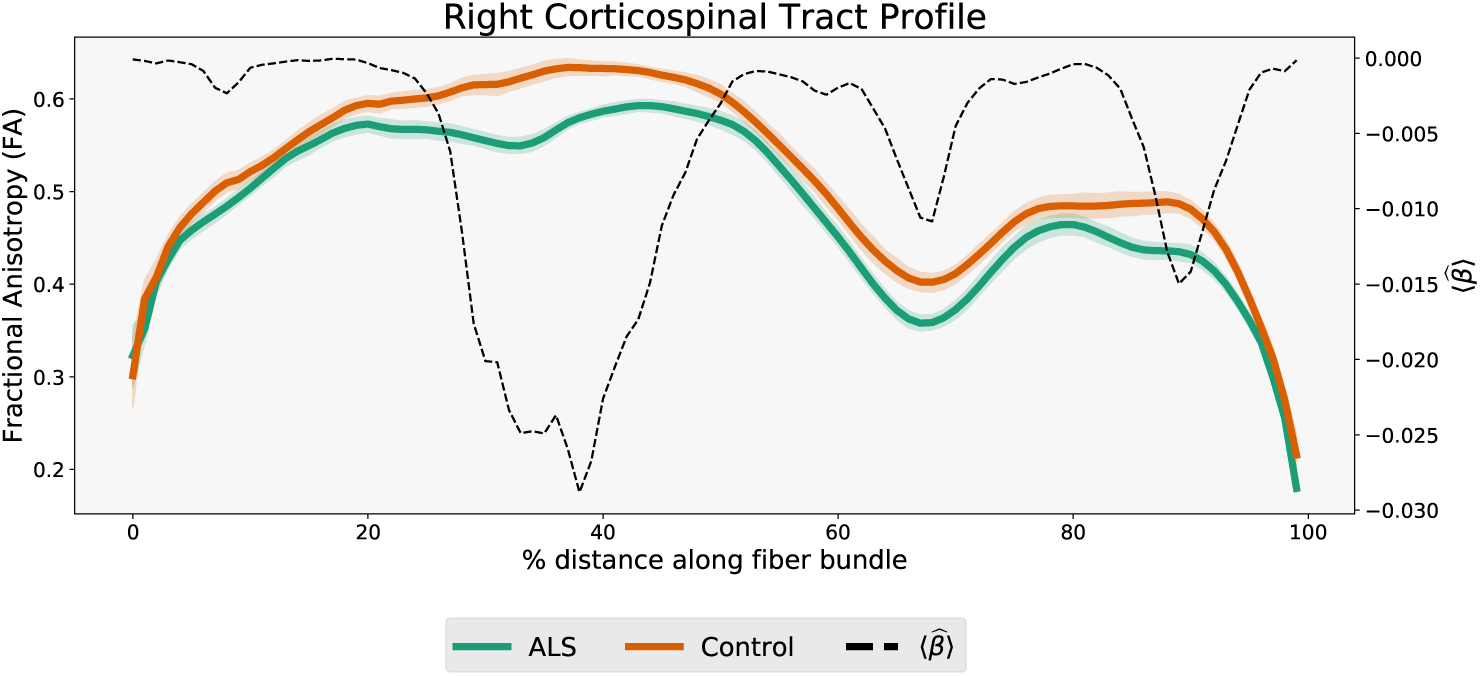
Model coefficients mirror FA differences. The places along the length of the CST where 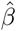 coefficients for FA (dashed line, right axis) have large negative values correspond to the locations of substantial differences between the ALS (green) and control (orange) FA (shaded area indicates standard error of the mean).

Analyzing the ways in which the model mislabels individuals may also provide insight. We found that mislabelled subjects are outliers relative to their group with respect to diffusion features of the CST. Figure 5 depicts the group FA values along with FA values of mislabelled subjects, two false negatives and one false-positive. The false negative classifications have high FA in one of the two sections of the CST where 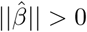 in Figure 4. The false positive subject has an FA that oscillates between the two group means. Thus, the SGL method fails comprehensibly.

**Fig 5.**
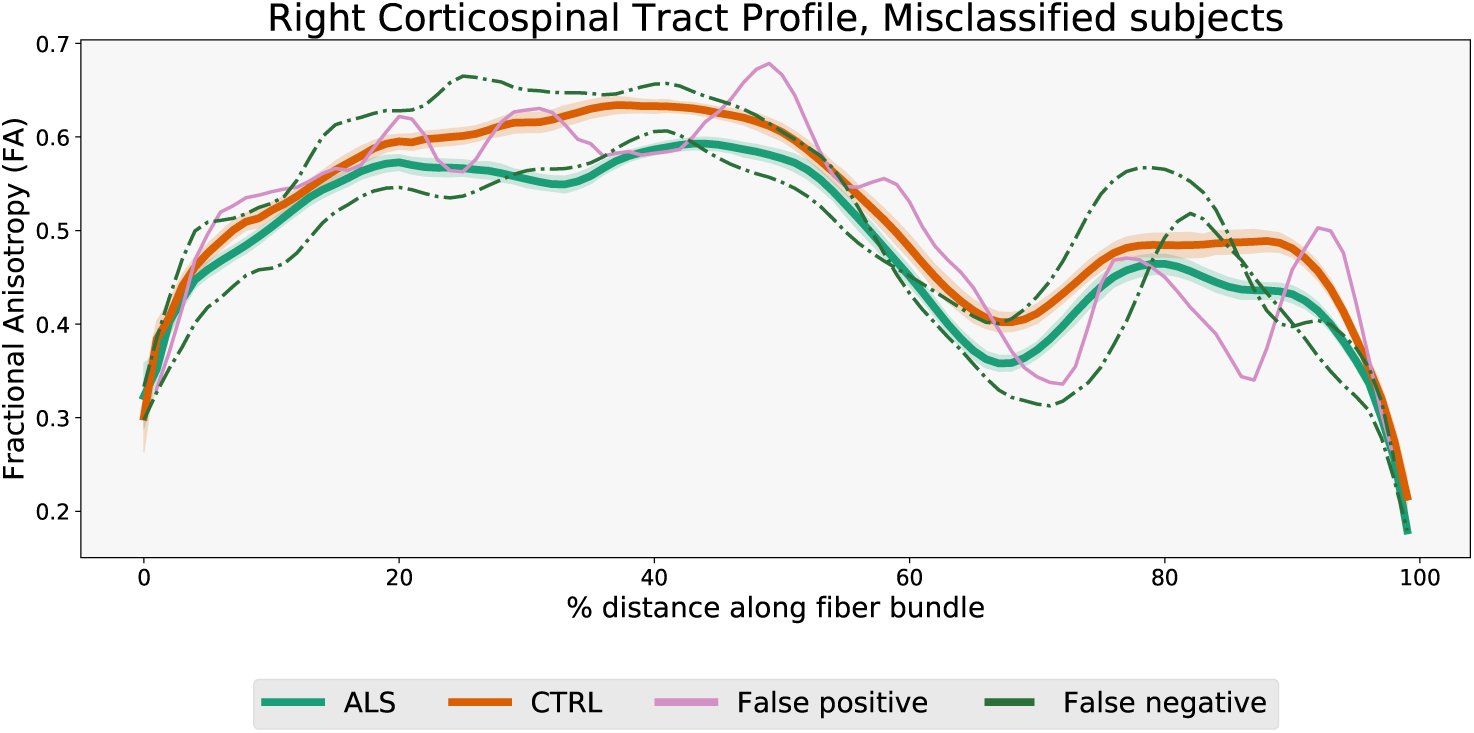
Model mis-classifications correspond to features identified by the model. The FA in CST of individuals that are mis-classified by the model is compared the group FA (with shaded are indicating standard error of the mean). False negative classifications (individuals that have ALS, but are classified as patients) correspond to high FA either in one of the two regions of large 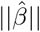 in Figure 4. The false positive classification has an FA profile that oscillates between the two group means.

### SGL accurately predicts age from tractometry data in a regression setting

To test the performance of SGL with tractometry data in a continuous regression task, we focus here on the prediction of biological age based on tractometry data. Prediction of “brain age” is a commonly undertaken task. This is both because it operates on a natural scale, with meaningful and easily understood units, as well as because predictions of brain age, and deviations from accurate prediction are diagnostic of overall brain health (for a recent review, see [61]). The WH dataset used here contains data from 76 healthy subjects, ranging between 6 years and 50 years of age [4]. In this case, biological age was used as the predicted variable (*y* in Eq (1)). SGL was fit to tractometry-extracted features: FA and MD in 20 major brain tracts, with each tract divided into 100 nodes. To evaluate the fit of the model, we used a nested cross-validation procedure. In this procedure, batches of subjects are held out. For each batch (or fold), the model is fully fit without this data. Then, once the parameters are fixed, the model is inverted to predict the ages of held out subjects based on the linear coeffiecients and the static non-linearity. This scheme automatically finds the right level of regularization (i.e., sparseness) and fits the coefficients to the ill-posed linear model, while guarding against overfitting. SGL accurately predicts the age of the subjects in this procedure, with a mean absolute error of 3.6 years (Figure 6, left panel). This is lower than the results of a recent study that predicted age in a large sample, based on diffusion MRI features [62]. Nevertheless, older subjects have higher residual variance, reflecting the automatically-chosen log-transformation and implying that brain age becomes more difficult to predict as we age chronologically (6, right panel). The model weights are distributed over many different tracts and dMRI tissue properties (Figure 7 left). This demonstrates that SGL is not coerced to produce overly sparse results when a more accurate model requires a dense selection of features. Furthermore, looking closer at a selection of tracts where high coefficients are found demonstrates that diffusion properties (FA, in this case) are different in different age groups in parts of the tracts where these higher coefficients are found (Figure 7 right).

**Fig 6.**
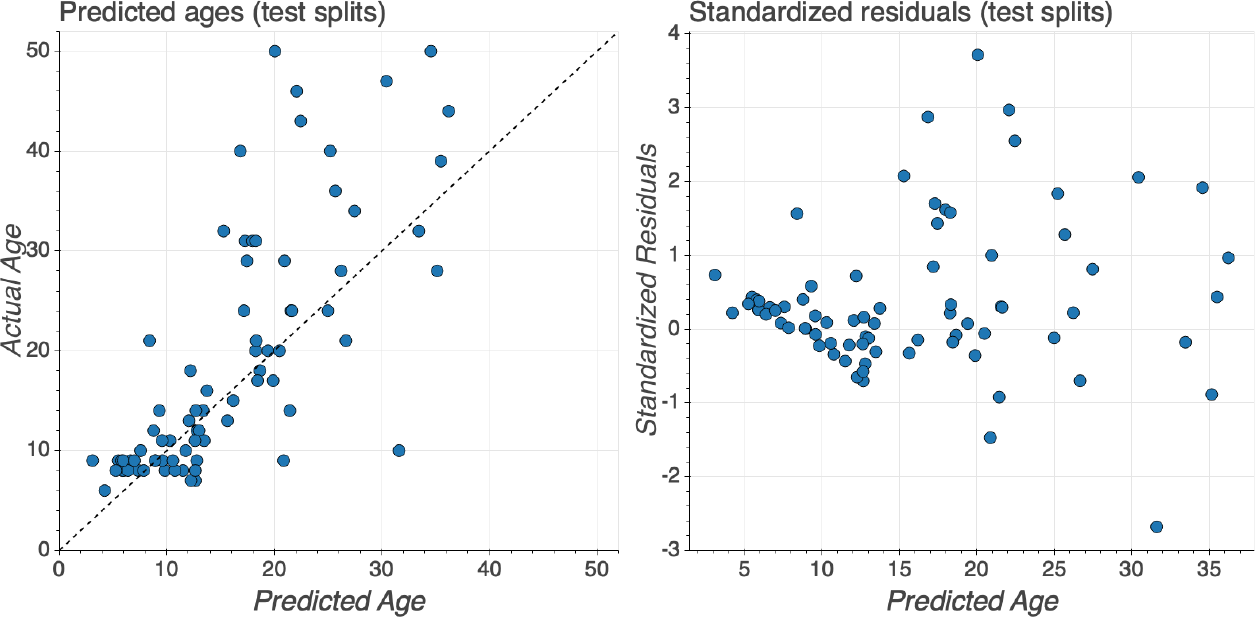
Predicting age with tractometry and SGL. Left: The predicted age of each individual (on the abscissa) and true age (on the ordinate), from the test splits (i.e., when each subject’s data was held out in fitting the model); an accurate prediction falls close to the *y* = *x* line (dashed). The mean absolute error in this case is 3.6 years and, the coefficient of determination *R*^2^ = 0.3. Right: Standardized residuals (on the abscissa) as a function of the true age (on the ordinate). Predictions are generally more accurate for younger individuals.

**Fig 7.**
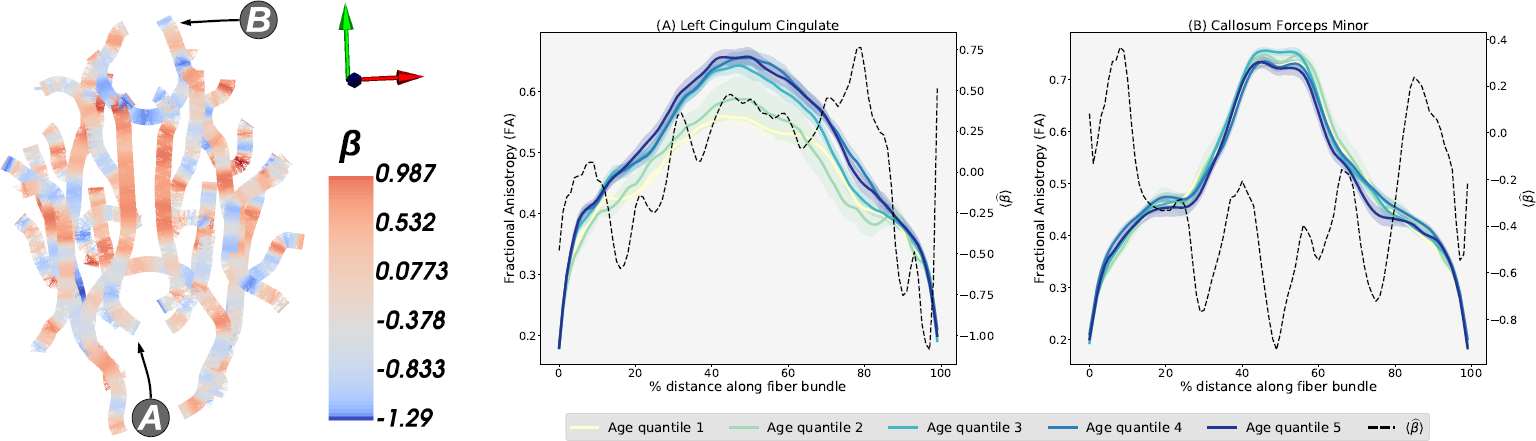
Feature importance for predicting age from tractometry. Left: A skeletonized display of the main brain tracts analyzed, with anterior facing up, and right hemisphere on the right. The 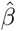 coefficients displayed in blue (negative) to red (positive) are for measurements of FA along the length of the tracts. The left cingulum cingulate (A) and forceps minor (B) are highlighted. Right: the FA (in shades of blue and green) and the *beta* coefficients (dashed) in (A) left cingulum and (B) forceps minor.

## Conclusion

We present here a novel method for analysis of dMRI tractometry data that relies on the Sparse Group Lasso [2] to (a) make accurate predictions of phenotypic properties of individual subjects while, simultaneously, (b) identifying the features of the white matter that are most important for this prediction in a completely data-driven approach. The method is broadly applicable to a wide range of research questions: it performs well in predicting both continuous variables, such as biological age, as well as categorical variables, such as whether a person is a patient or a healthy control. In both of these cases, SGL out-performs previous algorithms that have been developed for these tasks [3, 62]. The nested cross-validation approach used to fit the model and make both predictions and inferences from the model guards against overfitting and tunes the degree of sparseness required by the algorithm. This means that SGL can accurately describe phenomena that are locally confined to a particular anatomical location or diffusion property (e.g., FA in the CST) as well as phenomena that are widely distributed amongst brain regions and measured diffusion properties.

Specifically, we demonstrated that the algorithm correctly detects the fact that ALS, which is a disease of lower motor neurons, is localized to the cortico-spinal tract. This recapitulates the results of previous analysis of these same data, using a targeted ROI-based approach [3]. In contrast, for the analysis of biological age, the coefficients identified by the algorithm are very widely distributed across many parts of the white matter, mirroring previous results with this dataset (and others) that show a large and continuous distribution of life-span changes in white matter properties [4].

Taken together, these results demonstrate the promise of the group-regularized regression approach. Even at the scale of dozens of subjects, the results provided by SGL are both accurate, as well as interpretable [29]: tractometry capitalizes on domain knowledge to engineer meaningful features; SGL scores these features based on their relative importance; enables a visualization of these feature importance scores in the anatomical coordinate frame of the bundles (e.g., Figures 3 and fig:regress-beta) and provides a means to understand model errors (e.g., Figure 5). Thus, this multivariate analysis approach both (a) achieves high cross-validated accuracy for precision medicine applications of dMRI data and (b) identifies relevant features of brain anatomy that can further our neuroscientific understanding of clinical disorders.

Neuroscience has entered an era in which consortium efforts are putting together large datasets of high-quality dMRI measurements to address a variety of scientific questions [42, 63–66]. The volume and complexity of these data pose a substantial challenge. Dimensionality reduction with tractometry, followed by analysis with the approach we present here promises to capitalize on the wealth of data and on the co-measurement of interesting and important phenotypical data about brain health and about the participants’ cognitive abilities. We also expect the group-regularized approach to improve with larger datasets.

SGL has many other potential applications in neuroscience, because of the hierarchical and grouped nature of many data types that are collected in multiple sample points within anatomically-defined areas. For example, this method may be useful to understand the relationship between fMRI recordings and behavior, where activity in each voxel may co-vary with voxels within the same anatomical region and form features and groups of features. Similarly, large-scale multi-electrode recordings of neural activity in awake behaving animals are becoming increasingly feasible [67, 68] and these recordings can form features (neurons) and groups (neurons within an anatomical region). More ambitiously perhaps, this approach may be used to understand the role of correlations in so-called resting-state fMRI time-series and behavior, where pairwise correlations between voxels in different anatomical regions are features in the matrix and features may be grouped by pairs of anatomical regions. Given the large number of voxels in the surface of the gray matter and given that correlations increase the number of features by a factor of *n*^2^, this would pose a challenging problem to solve using SGL.

The results we present here also motivate extensions of the method using more sophisticated cost functions. For example, the fused sparse group lasso (FSGL) [69] extends SGL to enforce additional spatial structure: smoothness in the variation of diffusion metrics along the bundles. As brain measurements include additional structure (for example, bilateral symmetry), future work could also incorporate overlapping group membership for each entry in the tract profiles [70]. For example, a measurement could come from the corpus callosum, but also from the right hemipshere. This would also require extending the cost function used here to incorporate these constraints. Similarly, unsupervised dimensionality reduction of tractometry data (e.g., [71]) could also benefit from constraints based on grouping.

The method is packaged as open-source software called AFQ-Insight that is openly available, and provides a clear API to allow for extensions of the method. The sofware integrates within a broader automated fiber quantification software ecosystem: AFQ [1], which extracts tractometry data from raw and processed dMRI datasets, as well as AFQ-Browser, which visualizes tractometry data and facilitates sharing of the results of dMRI studies [49]. To facilitate reproducibility and ease use of the software, the results presented in this paper are also provided in https://github.com/richford/AFQ-Insight/tree/master/examples/preprint-notebooks as a series of Jupyter notebooks [72].

## Acknowledgments

This work was supported by BRAIN Iniative grant 1RF1MH121868-01 from the National Institutes for Mental Health and by a grant from the Gordon & Betty Moore Foundation and the Alfred P. Sloan Foundation to the University of Washington eScience Institute Data Science Environment. We would like to thank Scott Murray for a useful discussion of the SGL method and Mareike Grotheer for helpful comments on the manuscript.

## Notes

https://github.com/richford/afq-insight-paper

